# From Multi-Allele Fish to Non-Standard Environments, How ZFIN Assigns Phenotypes, Human Disease Models, and Gene Expression Annotations to Genes

**DOI:** 10.1101/2022.12.05.519159

**Authors:** Yvonne M. Bradford, Ceri E. Van Slyke, Douglas G. Howe, David Fashena, Ken Frazer, Ryan Martin, Holly Paddock, Christian Pich, Sridhar Ramachandran, Leyla Ruzicka, Amy Singer, Ryan Taylor, Wei-Chia Tseng, Monte Westerfield

**Affiliations:** The Institute of Neuroscience, University of Oregon, Eugene, Oregon 97403-1254, USA

## Abstract

*Danio rerio* is a model organism used to investigate vertebrate development. Manipulation of the zebrafish genome and resultant gene products by mutation or targeted knockdown has made the zebrafish a good system for investigating gene function, providing a resource to investigate genetic contributors to phenotype and human disease. Phenotypic outcomes can be the result of gene mutation, targeted knockdown of gene products, manipulation of experimental conditions, or any combination thereof. Zebrafish have been used in various genetic and chemical screens to identify genetic and environmental contributors to phenotype and disease outcomes. The Zebrafish Information Network (ZFIN) is the central repository for genetic, genomic, and phenotypic data that result from research using *Danio rerio*. Here we describe how ZFIN annotates phenotype, expression, and disease model data across various experimental designs, how we computationally determine wild-type gene expression, the phenotypic gene, and how these results allow us to propagate gene expression, phenotype, and disease model data to the correct gene, or gene related entity.

## Introduction

Understanding gene and protein function can provide insight to elucidate the intricate cellular mechanisms that are responsible for the development, growth, pathology, and senescence of organisms. Observing the results of gene mutation is the cornerstone of elucidating and understanding gene function. The zebrafish, *Danio rerio*, has been used in forward and reverse genetic screens to study gene function and understand the mechanisms of vertebrate development (Haffter *et al*. 1996)(Driever *et al*. 1996)(Moens *et al*. 2008)(Golling *et al*. 2002)(Varshney *et al*. 2013). The results of gene function studies in zebrafish are relevant to understanding human gene function due to the conservation of gene sequences and functions between zebrafish and humans (Howe *et al*. 2013a)(Postlethwait *et al*. 2000). Due to similarities between zebrafish and human organ functions and physiology, zebrafish have been used to model human diseases that affect the cardiovascular (Smith *et al*. 2009) (Liu *et al*. 2019), nervous (Chapman *et al*. 2013) (Hin *et al*. 2020), visual (Zhang *et al*. 2016), muscular (Majczenko *et al*. 2012) (Widrick *et al*. 2016), and many other systems. In addition to understanding gene function and disease modeling, zebrafish are increasingly used for toxicology and drug discovery studies, as well as research that explores the effects of genotype and environment on phenotype and disease (Zon and Peterson 2005) (Kaufman *et al*. 2009) (Cassar *et al*. 2019) (Williams *et al*. 2014)(Wheeler *et al*. 2019)

The Zebrafish Information Network, ZFIN, is the database resource for zebrafish research that annotates, curates, and makes data available from zebrafish research that spans genetic perturbations, chemically induced phenotypes, and human disease models, as well as gene expression (Sprague *et al*. 2008)(Ruzicka *et al*. 2015)(Howe *et al*. 2017). ZFIN curates gene expression, phenotype, and human disease model data by annotating the genotypes, experimental conditions, anatomical structures, phenotype statements, and disease models reported in zebrafish research publications (Sprague *et al*. 2006)(Howe *et al*. 2013b)(Bradford *et al*. 2017). These annotations can include genotypes with one or many alleles and experimental conditions that range from standard conditions to manipulation of temperature, diet, chemical, or other diverse conditions. Due to the breadth of data that represent combinations of genotype and environment that produce a phenotypic outcome or human disease model, it can be challenging to determine whether a particular allele or environment is causative. To understand gene function and clarify how gene dysfunction contributes to disease, it is necessary to separate genetic phenotypes from those caused by the environment. ZFIN has developed a data model and algorithms that distinguish the genotype and environment components of an annotation to parse genetic and environmental contributors to phenotypes, using the results to infer which genes are causative of a phenotype. Here we discuss the ZFIN annotation components and computational logic used to infer wild-type gene expression, gene-phenotype and gene-human disease relationships and the ZFIN webpages and download files where the data are available.

### ZFIN Annotation Components

There are three main components to ZFIN gene expression, phenotype, and human disease model annotations: 1) the genotype of the fish including gene knockdown reagents used (Fish), 2) the experimental conditions applied, and 3) an ontological representation of the results.

#### Fish

Zebrafish are an effective vertebrate model organism to understand gene function through gene mutation. Mutant gene loci are curated as alleles of genes and are part of a genotype together with the background strain when that information is provided. Zebrafish are also amenable to transgene insertion to knock out genes, insert mutant genes, or over-express genes to manipulate gene function (Amsterdam *et al*. 2004) (Kimelman *et al*. 2017) (Clark *et al*. 2011). ZFIN creates transgenic allele records for transgene insertions, and these alleles are represented in the genotype when applicable. Site specific mutagenesis using CRISPRs and TALENs is also used in zebrafish to screen for candidate genes (Jao *et al*. 2013) (Zu *et al*. 2013). Zebrafish crispants, F0 founder zebrafish created using CRISPRs, are also used to phenocopy loss of function mutants (Bek *et al*. 2021). In addition, gene function can be investigated in zebrafish using morpholinos, which knockdown the gene by targeting RNA, effectively silencing the gene product (Nasevicius and Ekker 2000) (Ekker and Larson 2001). ZFIN groups Morpholinos, CRISPRs, TALENs in a class called Sequence Targeting Reagents (STR) due to the sequence specific nature of these reagents. Both alleles and STRs have relationships to the genes they knockout or target. Because there are many ways in which gene function can be investigated in zebrafish, a flexible data model is needed to determine causative genes. To understand all of the genes that are affected due to either mutation or knockdown, ZFIN uses a data model that groups the genotype and applied STR in an object called Fish. The Fish represents the genotype and any STRs that have been used for gene knockdown.

#### Experimental Conditions

Zebrafish are used in a wide array of experimental contexts. To represent the experiments reported in research publications, the conditions applied are curated using ontology terms from the Zebrafish Experimental Conditions Ontology (ZECO) (Bradford *et al*. 2016) along with terms for the Zebrafish Anatomy Ontology (ZFA) (Van Slyke *et al*. 2014), Chemical Entities of Biological Interest (ChEBI) (Hastings *et al*. 2016), and NCBI Taxon (Federhen 2012). The ZECO ontology contains the main types of conditions with high-level nodes that include standard conditions for zebrafish husbandry as described in The Zebrafish Book (Westerfield 2000), control conditions (such as vehicle injections), biological treatment (such as exposure to bacteria), chemical treatment, diet alterations, housing conditions, *in vitro* culture, surgical manipulation, lighting conditions, temperature exposure, radiation exposure, and water quality. ZECO terms from the biological treatment branch are combined with NCBI Taxon terms to annotate conditions where another organism is added to the environment or when the zebrafish are raised in germ-free environments. The chemical treatment branch of ZECO is combined with chemicals from the ChEBI ontology to annotate the chemical that was used in the experiment. The surgical manipulation branch is combined with terms from the ZFA ontology to denote the anatomical structures that underwent ablation, resections, or other surgical manipulations. For instances when a cellular component, such as an axon, is ablated, GO-CC terms are used along with ZFA terms.

#### Ontological Representation of Results

ZFIN uses multiple ontologies to annotate gene expression, phenotype, human disease model, and gene function results. Disease, expression, and phenotype annotations include the Fish and experimental conditions. To complete disease annotations, terms from the Disease Ontology (DO) (Schriml *et al*. 2019) are added. To describe the location of the expression or phenotype annotation, terms from the ZFA, the Zebrafish Stage Ontology (ZFS) (Van Slyke *et al*. 2014), Gene Ontology Cellular Compartment (GO-CC)(Ashburner *et al*. 2000)(Carbon *et al*. 2019), Spatial Ontology (BSPO) (Dahdul *et al*. 2014), and ChEBI are used. Expression annotations include the gene that is expressed as well as the assay type using terms from the Measurement Method Ontology (MMO)(Smith *et al*. 2013). Phenotypes pertaining to the biological process or molecular function of a gene use GO Molecular Function (GO-MF) or GO Biological Process (GO_BP) terms in the annotation. All phenotype annotations use terms from the Phenotype and Trait Ontology (PATO) (Gkoutos *et al*. 2005). Gene function is annotated with the gene symbol and gene function terms from the Gene Ontology (GO) (Ashburner *et al*. 2000)(Carbon *et al*. 2019) as well as evidence terms from the Evidence and Conclusion Ontology (ECO)(Nadendla *et al*. 2022). All ZFIN annotations reference the publication that reported the results.

In summary, ZFIN gene expression, phenotype, and disease model annotations are multipartite including the genotype and applied knockdown reagents as Fish, the experimental conditions, and the ontological representation of the results. See Table 1a-c for examples of gene expression, phenotype, and human disease model annotations.

**Table 1a.** Gene Expression annotations

| Gene | Fish | Experimental Condition | Stage | Expression | Reference |
| --- | --- | --- | --- | --- | --- |
| *pax2a* | AB | Standard conditions [ZECO:0000103] | Pharyngula:Prim-25 [ZFS:0000031] | Optic furrow [ZFA:0005491] | ZDB-PUB-180407-9; PMID: 29625437 |
| *pax2a* | *aldh1a*<sup>i26/i26</sup> | Standard conditions [ZECO:0000103] | Segmentation:10-13 somites [ZFS:0000025] | Lateral plate mesoderm [ZFA:0000121] | ZDB-PUB-011002-4; PMID: 11688558 |
| *pax2a* | *cyp26a1*<sup>rw716/rw716</sup> | Chemical treatment: all-trans-retinoic acid [ZECO:0000111], [CHEBI:15367] | Segmentation: 1-4 somites [ZFS:0000023] | Midbrain hindbrain boundary neural keel [ZFA:0007045] | ZDB-PUB-061227-41; PMID: 17164423 |
| *pax2a* | AB+MO6-pax8+MO7-pax8 | Standard conditions [ZECO:0000103] | Segmentations: 5-9 somites [ZFS:0000024] | Epibranchial field [ZFA:0007061] | ZDB-PUB-110119-6; PMID: 21215261 |

**Table 1b.**
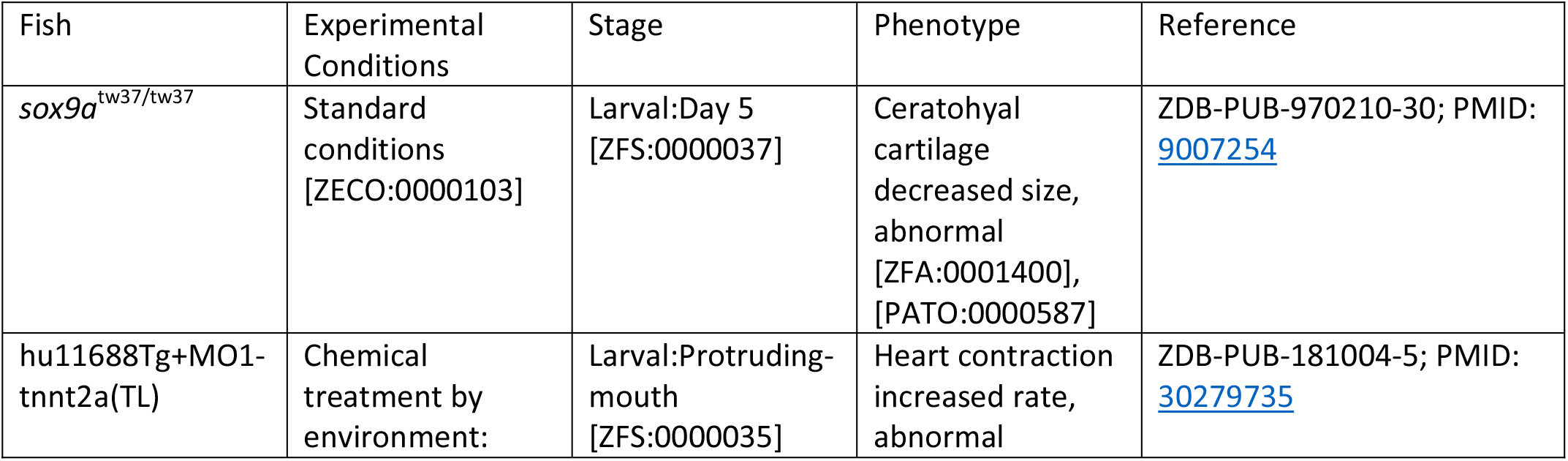
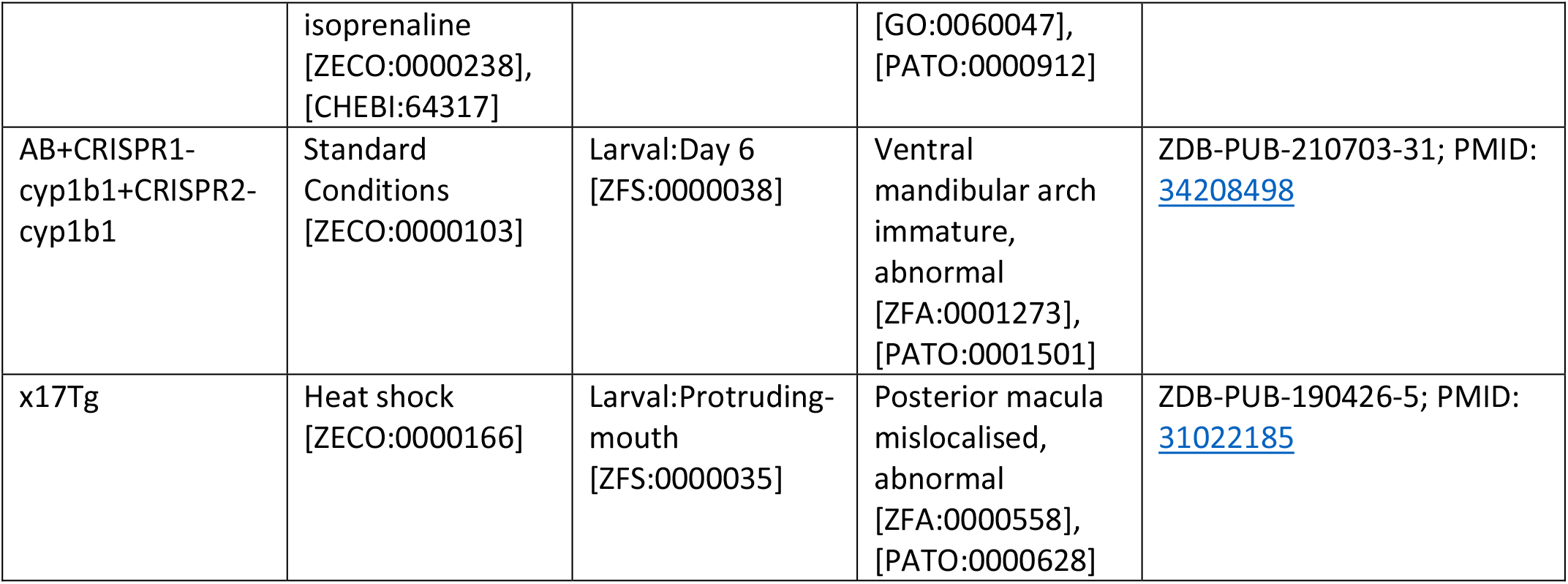
Phenotype annotations

| Fish | Experimental Conditions | Stage | Phenotype | Reference |
| --- | --- | --- | --- | --- |
| *sox9a*<sup>tw37/tw37</sup> | Standard conditions [ZECO:0000103] | Larval:Day 5 [ZFS:0000037] | Ceratohyal cartilage decreased size, abnormal [ZFA:0001400], [PATO:0000587] | ZDB-PUB-970210-30; PMID: 9007254 |
| hu11688Tg+MO1-tnnt2a(TL) | Chemical treatment by environment: | Larval:Protruding-mouth [ZFS:0000035] | Heart contraction increased rate, abnormal | ZDB-PUB-181004-5; PMID: 30279735 |
|  | isoprenaline<br>[ZECO:0000238],<br>[CHEBI:64317] |  | [GO:0060047],<br>[PATO:0000912] |  |
| AB+CRISPR1-<br>cyp1b1+CRISPR2-<br>cyp1b1 | Standard<br>Conditions<br>[ZECO:0000103] | Larval:Day 6<br>[ZFS:0000038] | Ventral<br>mandibular arch<br>immature,<br>abnormal<br>[ZFA:0001273],<br>[PATO:0001501] | ZDB-PUB-210703-31; PMID:<br>34208498 |
| x17Tg | Heat shock<br>[ZECO:0000166] | Larval:Protruding-mouth<br>[ZFS:0000035] | Posterior macula<br>mislocalised,<br>abnormal<br>[ZFA:0000558],<br>[PATO:0000628] | ZDB-PUB-190426-5; PMID:<br>31022185 |

**Table 1c.**
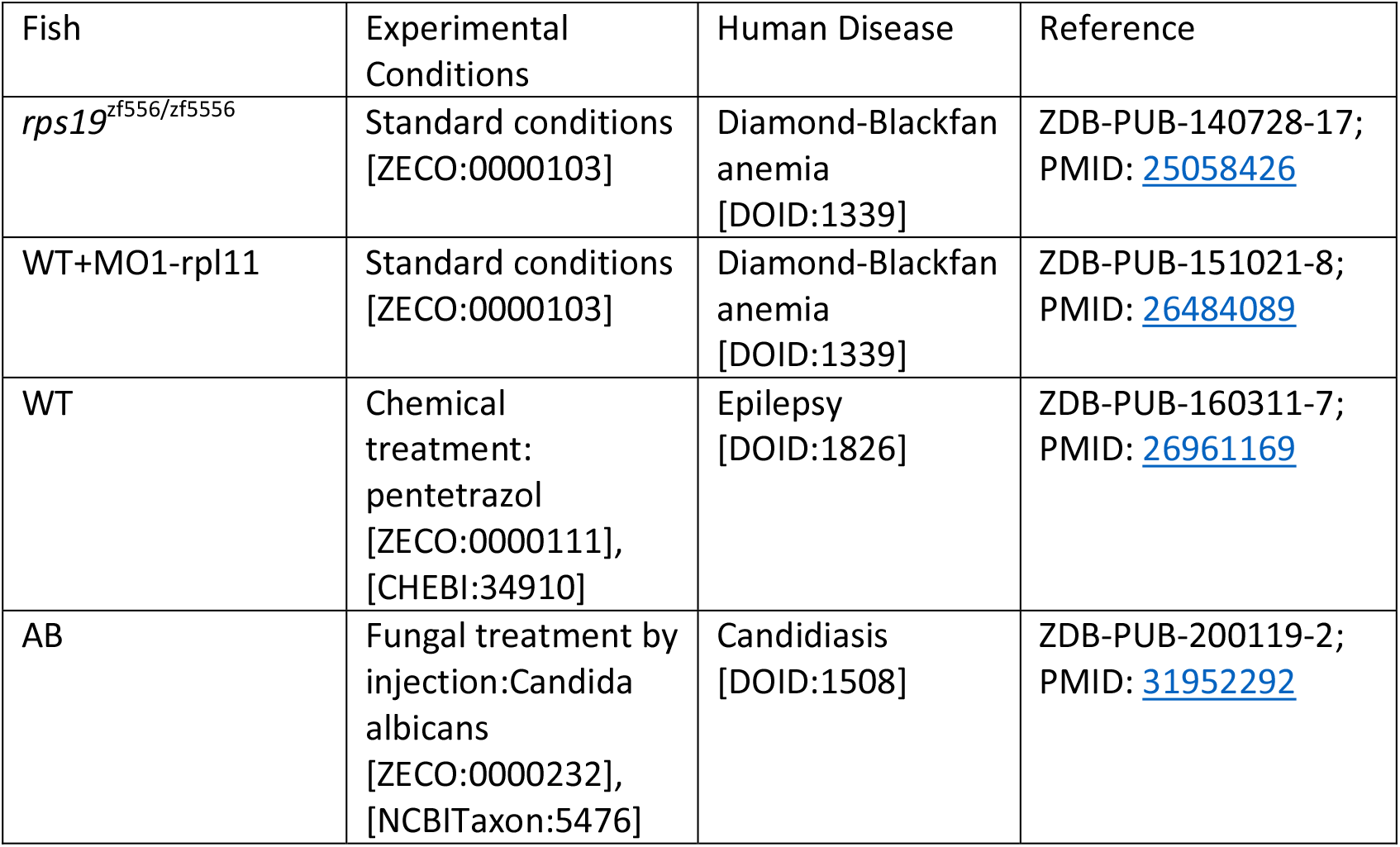
Human disease model annotations

| Fish | Experimental<br>Conditions | Human Disease | Reference |
| --- | --- | --- | --- |
| *rps19*<sup>zf556/zf5556</sup> | Standard conditions<br>[ZECO:0000103] | Diamond-Blackfan<br>anemia<br>[DOID:1339] | ZDB-PUB-140728-17;<br>PMID: 25058426 |
| WT+MO1-rpl11 | Standard conditions<br>[ZECO:0000103] | Diamond-Blackfan<br>anemia<br>[DOID:1339] | ZDB-PUB-151021-8;<br>PMID: 26484089 |
| WT | Chemical<br>treatment:<br>pentetrazol<br>[ZECO:0000111],<br>[CHEBI:34910] | Epilepsy<br>[DOID:1826] | ZDB-PUB-160311-7;<br>PMID: 26961169 |
| AB | Fungal treatment by<br>injection:Candida<br>albicans<br>[ZECO:0000232],<br>[NCBITaxon:5476] | Candidiasis<br>[DOID:1508] | ZDB-PUB-200119-2;<br>PMID: 31952292 |

### Database Logic for Gene Expression, Gene-Phenotype, Gene-Disease Associations

As described in the previous section, each data type provides different information used to construct an annotation. To be able to understand the function of a single gene, it is necessary to isolate the environmental factors from the genetic interactions within an annotation and ensure correct attribution of the experimental outcome to a single gene, if appropriate. To facilitate the correct representation of data sets and data displays, ZFIN has established query logic or algorithms to parse the details of existing annotations and display on the gene page those data that show where a gene is normally expressed and the phenotypic results of mutation or knockdown of that specific gene.

#### Wild-Type Gene Expression

Understanding the wild-type expression profile of genes is essential to understand what systems and structures a gene contributes to developmentally and is necessary as a comparator when evaluating gene expression in mutant or gene-knockdown zebrafish. ZFIN curators annotate gene expression in both wild-type and mutant backgrounds as well as what experimental conditions are present. To determine wild-type gene expression, algorithms are designed to identify gene expression in Fish that have wild-type backgrounds, no mutant alleles, in standard or control conditions. Gene expression results that meet these criteria are displayed on the gene page (Figure 1) and are provided in the ‘Expression data for wild-type fish’ download file available on the downloads page (https://zfin.org/downloads). Mutant or non-wild-type zebrafish gene expression can be found on the Fish page, via the search interface, in the download file ‘ZFIN Genes with Expression Assay Records’, and on STR pages. The STR page displays expression in Fish only where a single STR is used in a wild-type background (Figure 4).

**Figure 1.**
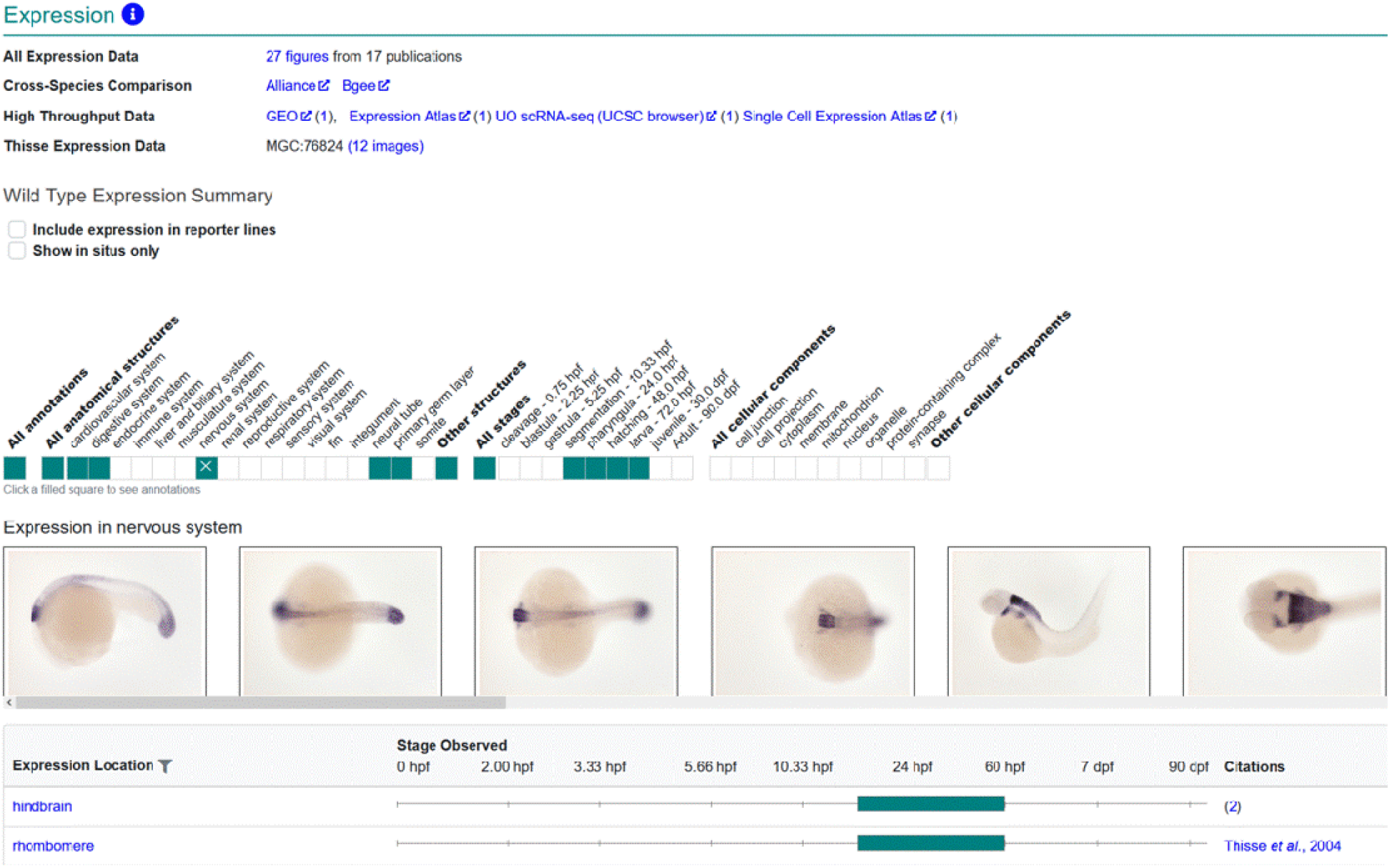
Gene page gene expression. Gene expression displayed on the gene page is limited to gene expression results in wild-type backgrounds. The Wild-Type Expression Summary displays a graphical ribbon that denotes the anatomical systems and stages that have gene expression annotations. The table lists the anatomical terms, stages and citations.

#### Affected Gene for Phenotype and Disease Model

To determine the function of a gene, it is instructive to look at the phenotypic outcomes of mutant and gene-knockdown zebrafish. Phenotype can encompass many levels of observation from morphologic changes at the level of the whole organism to changes in gene expression and protein location within a cell. To draw conclusions about what functions a gene has in the cell or organism, it is necessary to ensure that the phenotypes attributed to the gene are solely caused by changes to that gene. ZFIN has developed algorithms to determine the total number of altered or affected genes in a Fish, which is used to determine the causative gene. The number of affected genes is determined by counting distinct genes associated with alleles and STRs that are associated with a Fish. When the affected gene count equals one and the experimental conditions are standard/generic control, the phenotype or disease association is inferred or calculated to be caused by the gene associated to the Fish either by is_allele relationship or by STR target relationship. There are various ways to arrive at gene count = 1. As illustrated in Figure 2, Fish can have one affected gene but can be more or less complex in their genetic makeup. For example, a Fish with a single allele with one affected gene, a Fish with multiple alleles where all alleles affect the same gene, a wild-type Fish injected with one or more STRs targeting one gene, and a non-phenotypic transgenic line injected with one or more STRs that target one gene all have only a single affected gene.

**Figure 2.**
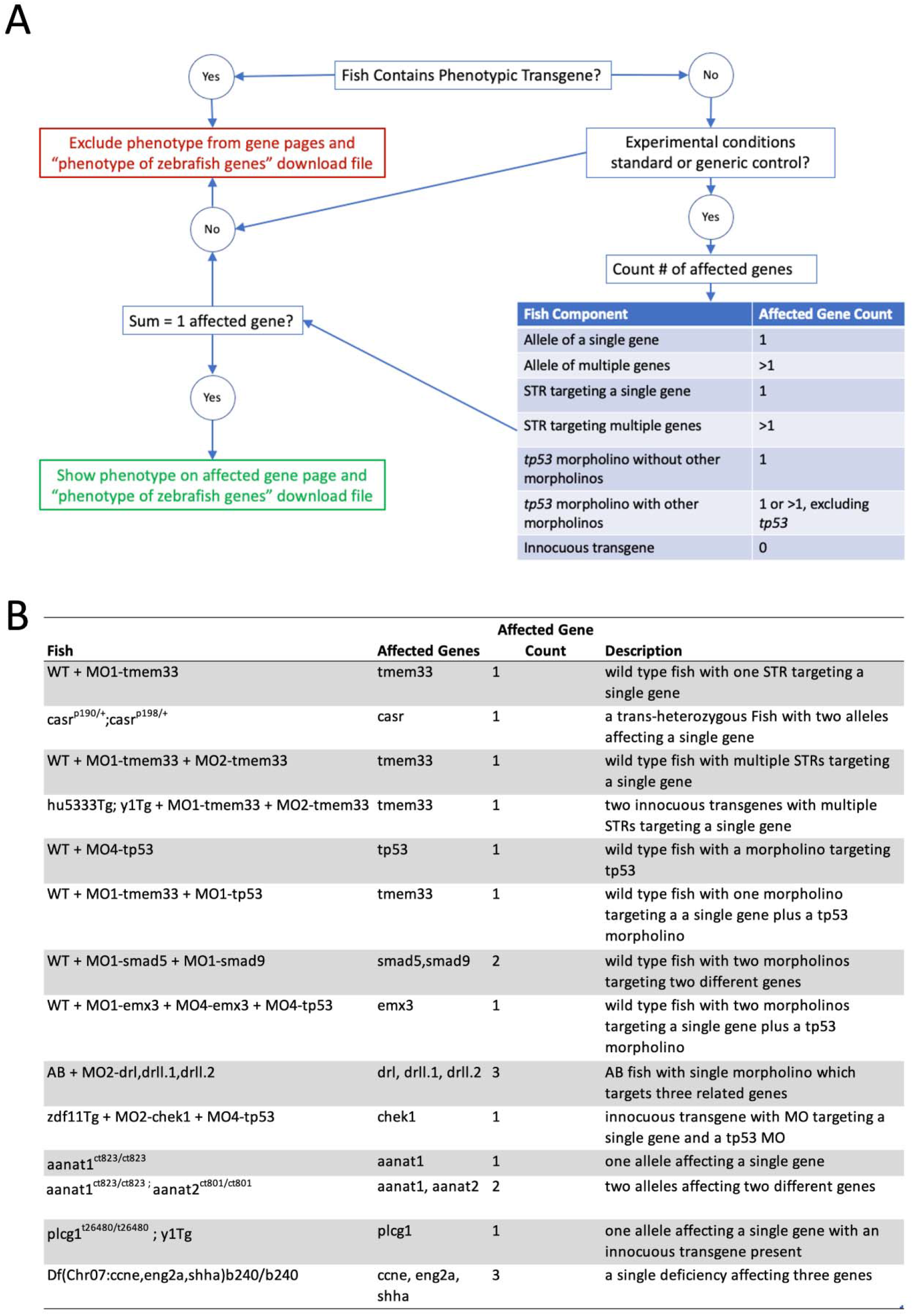
Logic for determining how many genes are affected in a Fish and whether or not associated phenotype data can be shown on a gene page. A) A logic flow diagram describing the algorithm used to determine how many genes are affected in a Fish and whether phenotype data can be shown on a gene page. B) A table of examples of Fish that result in variable numbers of affected genes.

We have recently added rules to the algorithm that do not count *tp53* as an affected gene in Fish where morpholinos against *tp53* were used in addition to other non-tp53 morpholinos. This rule accommodates the way zebrafish researchers use morpholinos against *tp53* to deal with non-specific effects of morpholinos (Robu *et al*. 2007). Previously, a Fish that had two morpholinos, one of which was against *tp53*, would be considered to have two affected genes and the phenotype would be excluded from gene pages. The algorithm now ignores *tp53* morpholinos in the Fish and the resulting group of morpholinos is used to obtain the affected gene count, with data propagated to the gene page when the gene count equals one (Figure 3).

**Figure 3.**
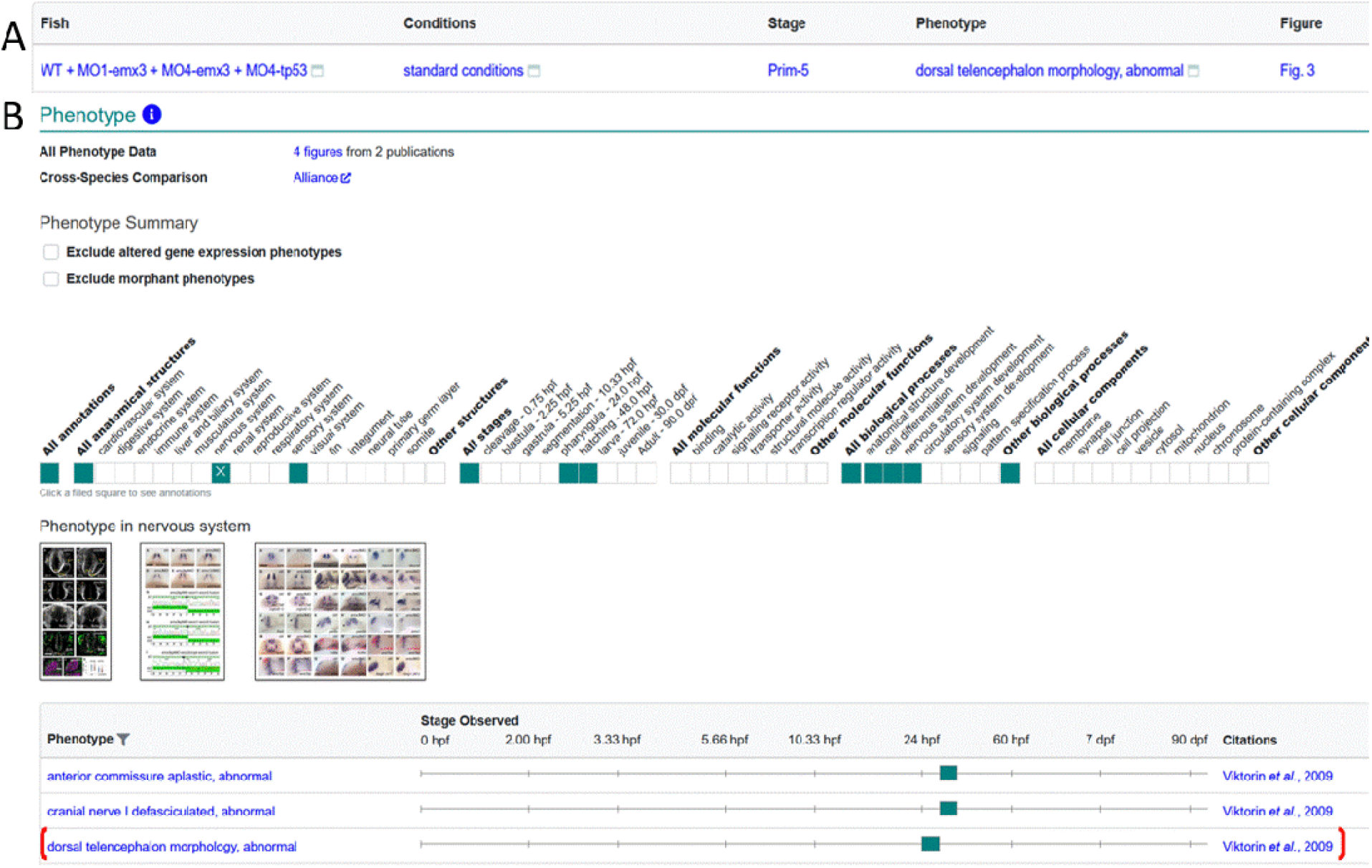
Display of MO-*tp53* Fish data on Gene page. A) Phenotype data for Fish WT+MO1-emx3+MO4-emx3+MO4-tp53 in standard conditions as reported in Viktorin *et al*. 2009. B) The phenotype summary section on the *emx3* gene page has a ribbon that denotes systems, stages, biological processes and cellular components that have annotations, with individual annotations displayed in the table. Thumbnail images are displayed when available. Phenotype corresponding to Fish in A is denoted by red bracket.

In addition to counting the number of affected genes the algorithms account for transgenic lines, both those that are treated as wild-type equivalents by the research community and those used to alter the expression of a gene. ZFIN annotates transgenic genomic features (alleles) as phenotypic or innocuous based on the transgenic construct associated with them. The phenotypic relationship is used with constructs that drive expression of either an endogenous zebrafish gene or a gene from another species (Table 2). These constructs are expected to produce protein products that can have a phenotypic effect. The innocuous relationship is used with constructs that drive the expression of fluorescent proteins, or are unable to transcribe a protein product unless inserted near a native promoter, such as gene trap constructs. Information on the innocuous or phenotypic relationship between a genomic feature and a construct is available in the ‘Innocuous/Phenotypic Construct Details’ download file. Fish containing genomic features that have a phenotypic relationship to a construct are excluded by affected gene count algorithms because phenotype and disease annotations using such Fish cannot be attributed to a single gene. Fish that have genomic features with an innocuous relationship to a construct are counted as wild-type equivalents by the affected gene count algorithms. The resulting data allow us to determine computationally the affected gene count. In addition to gene count and innocuous or transgenic genomic features, the experimental conditions are also taken into account when determining whether the phenotype or disease model data can be attributed to a gene. When the experimental conditions are standard or generic control and the affected gene count is one, the resulting phenotype or disease association is inferred to be caused by the one affected gene. These data are then propagated to the gene page, gene related entity pages, and download files.

**Table 2.**
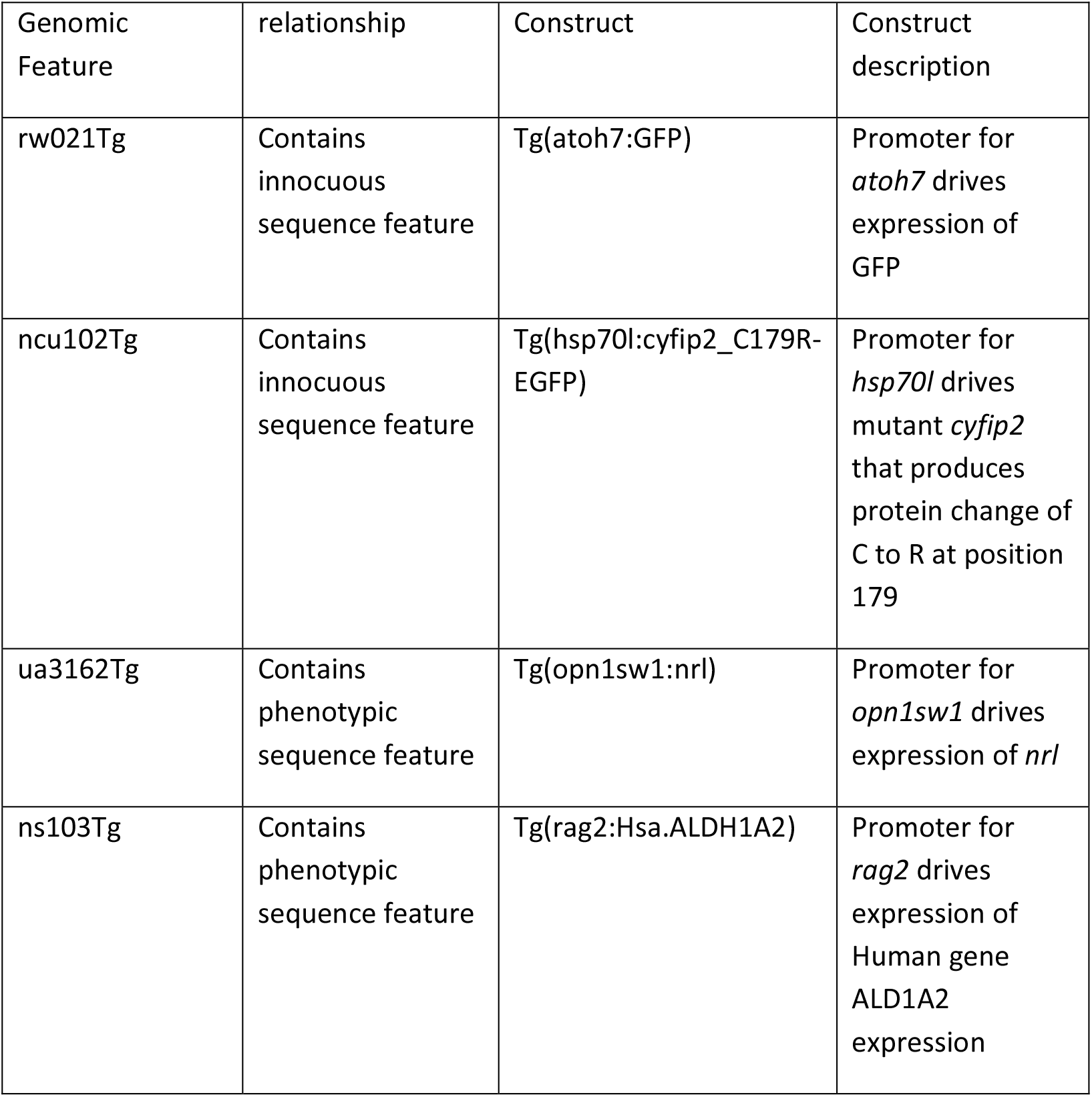
Innocuous and phenotypic constructs

Similar rules are employed for determining whether phenotype is caused by an STR or may be the result of a combination of genetic affectors. On the STR page, phenotype in Fish with only a single STR targeting a single gene in a wild-type or non-phenotypic transgenic background is displayed in the section where the label starts with “Phenotype resulting from” followed by the STR name (Figure 4). For more complex Fish or when the STR has multiple targets, the phenotypes are displayed in a section labeled “Phenotype of all Fish created by or utilizing” followed by the STR name(s).

**Figure 4.**
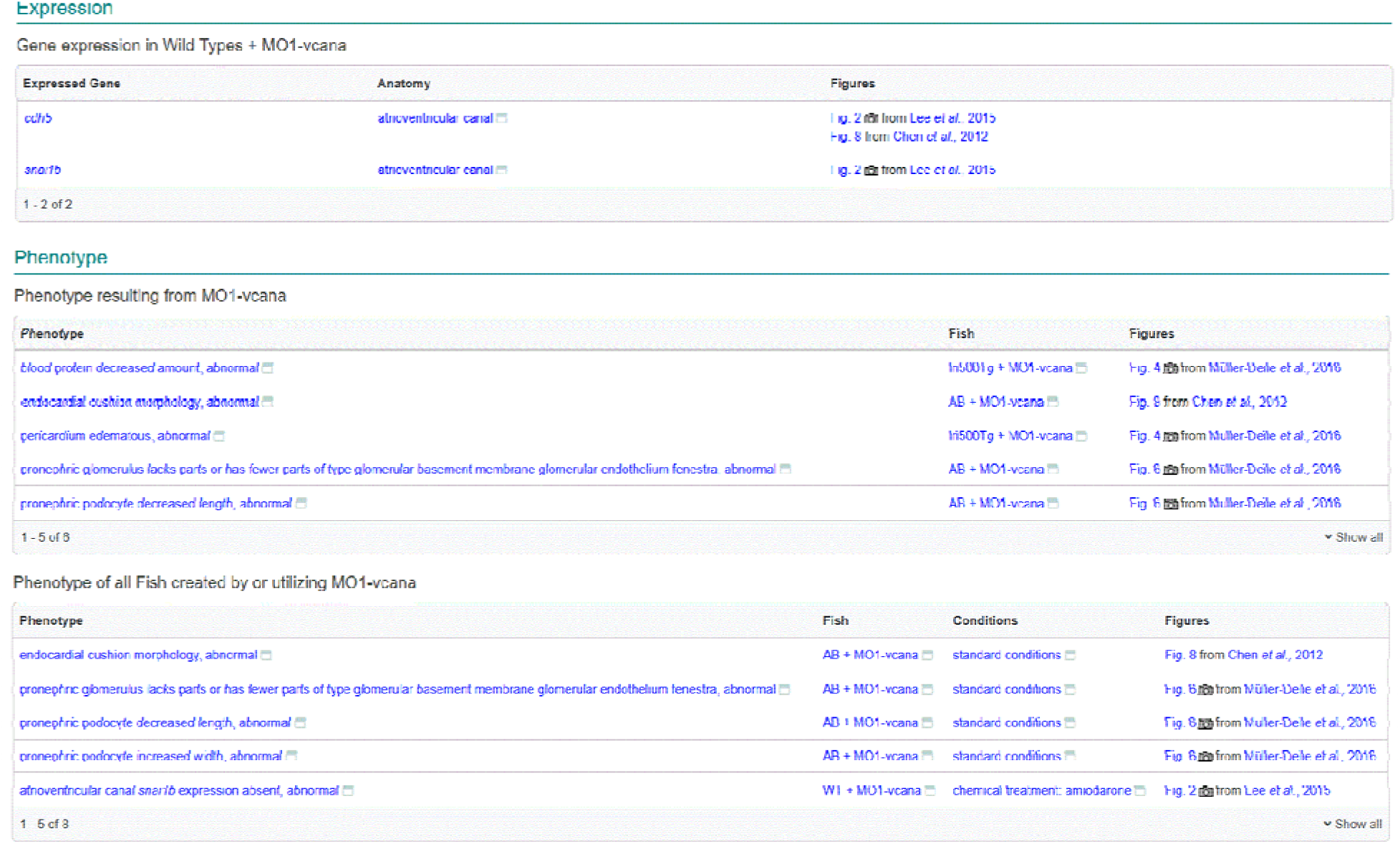
STR page. Expression display is limited to fish with a wild-type background under standard or control conditions. Phenotype display is divided into two sections, the first labeled “Phenotype resulting from MO1-vcana” contains phenotype only in wild-type or innocuous transgenic fish with standard conditions. Phenotype in more complex fish or under non standard conditions as well as the phenotype from the previous section is displayed in the section labeled “Phenotype of all Fish created by or utilizing MO1-vcana”.

## Conclusion

The development, growth and senescence of organisms is the result of an elegant orchestra of gene expression, protein function, pathology, and the environment. Understanding gene and protein function is essential knowledge that provides insight into the cellular mechanisms of developmental and disease processes. Gene function has traditionally been elucidated using gene mutation and targeted gene knockdown. Genetic and experimental condition manipulation, either singly or in combination, produces phenotypic outcomes. Zebrafish have been used in forward and reverse genetic screens to study gene function, model human disease, understand toxicology, and discover drugs. ZFIN curates genetic, genomic, phenotypic, and disease model data that result from zebrafish research. The algorithms used by ZFIN support the identification of wild-type expression patterns, genes that are causative for phenotypes, and disease models from data collected in a wide variety of genetic backgrounds and experimental conditions. The resulting data are presented on the gene, STR, and disease pages as well as in specialized download files. The aggregation of these data on discrete pages and download files allows users to quickly synthesize data about gene function, phenotypic outcomes and disease models without having to compile manually the research from many genotypes, gene knockdowns, and experimental conditions.

## References

Amsterdam A., R. M. Nissen, Z. Sun, E. C. Swindell, S. Farrington, et al., 2004 Identification of 315 genes essential for early zebrafish development. Proc. Natl. Acad. Sci. U. S. A. 101: 12792–12797. https://doi.org/10.1073/PNAS.0403929101

Ashburner M., C. A. Ball, J. A. Blake, D. Botstein, H. Butler, et al., 2000 Gene ontology: Tool for the unification of biology. Nat. Genet. 25: 25–29.

Bek J. W., C. Shochat, A. De Clercq, H. De Saffel, A. Boel, et al., 2021 Lrp5 Mutant and Crispant Zebrafish Faithfully Model Human Osteoporosis, Establishing the Zebrafish as a Platform for CRISPR-Based Functional Screening of Osteoporosis Candidate Genes. J. Bone Miner. Res. 36: 1749–1764. https://doi.org/10.1002/JBMR.4327

Bradford Y. M., C. E. Van Slyke, S. Toro, and S. Ramachandran, 2016 The zebrafish experimental conditions ontology: Systemizing experimental descriptions in ZFIN, in CEUR Workshop Proceedings,.

Bradford Y. M., S. Toro, S. Ramachandran, L. Ruzicka, D. G. Howe, et al., 2017 Zebrafish models of human disease: Gaining insight into human disease at ZFIN. ILAR J. 58: 4–16. https://doi.org/10.1093/ilar/ilw040

Carbon S., E. Douglass, N. Dunn, B. Good, N. L. Harris, et al., 2019 The Gene Ontology Resource: 20 years and still GOing strong. Nucleic Acids Res. 47: D330–D338. https://doi.org/10.1093/nar/gky1055

Cassar S., I. Adatto, J. L. Freeman, J. T. Gamse, aki Iturria, et al., 2019 Use of Zebrafish in Drug Discovery Toxicology. https://doi.org/10.1021/acs.chemrestox.9b00335

Chapman A. L., E. J. Bennett, T. M. Ramesh, K. J. De Vos, and A. J. Grierson, 2013 Axonal Transport Defects in a Mitofusin 2 Loss of Function Model of Charcot-Marie-Tooth Disease in Zebrafish. PLoS One 8. https://doi.org/10.1371/JOURNAL.PONE.0067276

Clark B. S., M. Winter, A. R. Cohen, and B. A. Link, 2011 Generation of Rab-based transgenic lines for in vivo studies of endosome biology in zebrafish. Dev. Dyn. 240: 2452–2465. https://doi.org/10.1002/DVDY.22758

Dahdul W. M., H. Cui, P. M. Mabee, C. J. Mungall, D. Osumi-Sutherland, et al., 2014 Nose to tail, roots to shoots: Spatial descriptors for phenotypic diversity in the Biological Spatial Ontology. J. Biomed. Semantics 5: 1–13. https://doi.org/10.1186/2041-1480-5-34

Driever W., L. Solnica-Krezel, A. F. Schier, S. C. F. Neuhauss, J. Malicki, et al., 1996 A genetic screen for mutations affecting embryogenesis in zebrafish. Development 123: 37–46. https://doi.org/10.1242/DEV.123.1.37

Ekker S. C., and J. D. Larson, 2001 Morphant technology in model developmental systems. Genesis 30: 89–93. https://doi.org/10.1002/GENE.1038

Federhen S., 2012 The NCBI Taxonomy database. Nucleic Acids Res. 40: D136–D143. https://doi.org/10.1093/NAR/GKR1178

Gkoutos G. V, E. C. J. Green, A. M. Mallon, J. M. Hancock, and D. Davidson, 2005 Using ontologies to describe mouse phenotypes. Genome Biol. 6. https://doi.org/10.1186/gb-2004-6-1-r8

Golling G., A. Amsterdam, Z. Sun, M. Antonelli, E. Maldonado, et al., 2002 Insertional mutagenesis in zebrafish rapidly identifies genes essential for early vertebrate development. Nat. Genet. 2002 312 31: 135–140. https://doi.org/10.1038/ng896

Haffter P., M. Granato, M. Brand, M. C. Mullins, M. Hammerschmidt, et al., 1996 The identification of genes with unique and essential functions in the development of the zebrafish, Danio rerio. Development 123: 1–36. https://doi.org/10.1242/DEV.123.1.1

Hastings J., G. Owen, A. Dekker, M. Ennis, N. Kale, et al., 2016 ChEBI in 2016: Improved services and an expanding collection of metabolites. Nucleic Acids Res. 44: D1214–D1219. https://doi.org/10.1093/nar/gkv1031

Hin N., M. Newman, J. Kaslin, A. M. Douek, A. Lumsden, et al., 2020 Accelerated brain aging towards transcriptional inversion in a zebrafish model of the K115fs mutation of human PSEN2. PLoS One 15. https://doi.org/10.1371/JOURNAL.PONE.0227258

Howe K., M. D. Clark, C. F. Torroja, J. Torrance, C. Berthelot, et al., 2013a The zebrafish reference genome sequence and its relationship to the human genome. Nature. https://doi.org/10.1038/nature12111

Howe D. G., Y. M. Bradford, T. Conlin, A. E. Eagle, D. Fashena, et al., 2013b ZFIN, the Zebrafish Model Organism Database: increased support for mutants and transgenics. Nucleic Acids Res. 41: D854–60. https://doi.org/10.1093/nar/gks938

Howe D. G. D. G., Y. M. Y. M. Bradford, A. Eagle, D. Fashena, K. Frazer, et al., 2017 The Zebrafish Model Organism Database: new support for human disease models, mutation details, gene expression phenotypes and searching. Nucleic Acids Res. 45: D758–D768. https://doi.org/10.1093/nar/gkw1116

Jao L. E., S. R. Wente, and W. Chen, 2013 Efficient multiplex biallelic zebrafish genome editing using a CRISPR nuclease system. Proc. Natl. Acad. Sci. U. S. A. 110: 13904–13909. https://doi.org/10.1073/PNAS.1308335110/-/DCSUPPLEMENTAL

Kaufman C. K., R. M. White, and L. Zon, 2009 Chemical Genetic Screening in the Zebrafish Embryo. Nat. Protoc. 4: 1422. https://doi.org/10.1038/NPROT.2009.144

Kimelman D., N. L. Smith, J. K. H. Lai, and D. Y. R. Stainier, 2017 Regulation of posterior body and epidermal morphogenesis in zebrafish by localized Yap1 and Wwtr1. Elife 6. https://doi.org/10.7554/ELIFE.31065

Liu L., F. Fei, R. Zhang, F. Wu, Q. Yang, et al., 2019 Combinatorial genetic replenishments in myocardial and outflow tract tissues restore heart function in tnnt2 mutant zebrafish. Biol. Open 8. https://doi.org/10.1242/BIO.046474

Majczenko K., A. E. Davidson, S. Camelo-Piragua, P. B. Agrawal, R. A. Manfready, et al., 2012 Dominant mutation of CCDC78 in a unique congenital myopathy with prominent internal nuclei and atypical cores. Am. J. Hum. Genet. 91: 365–371. https://doi.org/10.1016/J.AJHG.2012.06.012

Moens C. B., T. M. Donn, E. R. Wolf-Saxon, and T. P. Ma, 2008 Reverse genetics in zebrafish by TILLING. Briefings Funct. Genomics Proteomics 7: 454. https://doi.org/10.1093/BFGP/ELN046

Nadendla S., R. Jackson, J. Munro, F. Quaglia, B. Mészáros, et al., 2022 ECO: the Evidence and Conclusion Ontology, an update for 2022. Nucleic Acids Res. 50: D1515–D1521. https://doi.org/10.1093/NAR/GKAB1025

Nasevicius A., and S. C. Ekker, 2000 Effective targeted gene “knockdown” in zebrafish. Nat. Genet. 26: 216–20. https://doi.org/10.1038/79951

Postlethwait J. H., I. G. Woods, P. Ngo-Hazelett, Y. L. Yan, P. D. Kelly, et al., 2000 Zebrafish Comparative Genomics and the Origins of Vertebrate Chromosomes. Genome Res. 10: 1890–1902. https://doi.org/10.1101/GR.164800

Robu M. E., J. D. Larson, A. Nasevicius, S. Beiraghi, C. Brenner, et al., 2007 p53 activation by knockdown technologies. PLoS Genet. 3: 787–801. https://doi.org/10.1371/JOURNAL.PGEN.0030078

Ruzicka L., Y. M. Bradford, K. Frazer, D. G. Howe, H. Paddock, et al., 2015 ZFIN, The zebrafish model organism database: Updates and new directions. Genesis 53. https://doi.org/10.1002/dvg.22868

Schriml L. M., E. Mitraka, J. Munro, B. Tauber, M. Schor, et al., 2019 Human Disease Ontology 2018 update: Classification, content and workflow expansion. Nucleic Acids Res. 47: D955–D962. https://doi.org/10.1093/nar/gky1032

Slyke C. E. C. E. Van, Y. M. Y. M. Bradford, M. Westerfield, and M. A. M. A. Haendel, 2014 The zebrafish anatomy and stage ontologies: representing the anatomy and development of Danio rerio. J. Biomed. Semant. 5: 12. https://doi.org/10.1186/2041-1480-5-12

Smith K. A., I. C. Joziasse, S. Chocron, M. Van Dinther, V. Guryev, et al., 2009 Dominant-negative alk2 allele associates with congenital heart defects. Circulation 119: 3062–3069. https://doi.org/10.1161/CIRCULATIONAHA.108.843714/FORMAT/EPUB

Smith J. R., C. A. Park, R. Nigam, S. J. F. Laulederkind, G. T. Hayman, et al., 2013 The clinical measurement, measurement method and experimental condition ontologies: expansion, improvements and new applications. J. Biomed. Semantics 4. https://doi.org/10.1186/2041-1480-4-26

Sprague J., L. Bayraktaroglu, D. Clements, T. Conlin, D. Fashena, et al., 2006 The Zebrafish Information Network: the zebrafish model organism database. Nucleic Acids Res. 34: 581–585. https://doi.org/10.1093/nar/gkj086

Sprague J., L. Bayraktaroglu, Y. Bradford, T. Conlin, N. Dunn, et al., 2008 The Zebrafish Information Network: the zebrafish model organism database provides expanded support for genotypes and phenotypes. Nucleic Acids Res. 36: D768–72. https://doi.org/10.1093/nar/gkm956

Varshney G. K., J. Lu, D. E. Gildea, H. Huang, W. Pei, et al., 2013 A large-scale zebrafish gene knockout resource for the genome-wide study of gene function. Genome Res. 23: 727–735. https://doi.org/10.1101/GR.151464.112

Viktorin G., C. Chiuchitu, M. Rissler, Z. M. Varga, and M. Westerfield, 2009 Emx3 is required for the differentiation of dorsal telencephalic neurons. Dev. Dyn. 238: 1984–1998. https://doi.org/10.1002/DVDY.22031

Westerfield M., 2000 The zebrafish book: a guide for the laboratory use of zebrafish (Danio rerio). University of Oregon Press, Eugene, OR.

Wheeler M. A., M. Jaronen, R. Covacu, S. E. J. Zandee, G. Scalisi, et al., 2019 Environmental Control of Astrocyte Pathogenic Activities in CNS Inflammation. Cell 176: 581-596.e18. https://doi.org/10.1016/J.CELL.2018.12.012

Widrick J. J., M. S. Alexander, B. Sanchez, D. E. Gibbs, G. Kawahara, et al., 2016 Muscle dysfunction in a zebrafish model of Duchenne muscular dystrophy. Physiol. Genomics 48: 850–860. https://doi.org/10.1152/PHYSIOLGENOMICS.00088.2016

Williams T. D., L. Mirbahai, and J. K. Chipman, 2014 The toxicological application of transcriptomics and epigenomics in zebrafish and other teleosts. Brief. Funct. Genomics 13: 157–171. https://doi.org/10.1093/BFGP/ELT053

Zhang J., C. Wang, Y. Shen, N. Chen, L. Wang, et al., 2016 A mutation in ADIPOR1 causes nonsyndromic autosomal dominant retinitis pigmentosa. Hum Genet 135: 1375–1387. https://doi.org/10.1007/s00439-016-1730-2

Zon L. I., and R. T. Peterson, 2005 In vivo drug discovery in the zebrafish. Nat. Rev. Drug Discov. 2005 41 4: 35–44. https://doi.org/10.1038/nrd1606

Zu Y., X. Tong, Z. Wang, D. Liu, R. Pan, et al., 2013 TALEN-mediated precise genome modification by homologous recombination in zebrafish. Nat. Methods 2013 104 10: 329–331. https://doi.org/10.1038/nmeth.2374

